# Howler monkeys are the reservoir of malaria parasites causing zoonotic infections in the Atlantic forest of Rio de Janeiro

**DOI:** 10.1101/715623

**Authors:** Filipe Vieira Santos de Abreu, Edmilson dos Santos, Aline Rosa Lavigne Mello, Larissa Rodrigues Gomes, Denise Anete Madureira de Alvarenga, Marcelo Quintela Gomes, Waldemir Paixão Vargas, Cesare Bianco-Júnior, Anielle de Pina-Costa, Danilo Simonini Teixeira, Alessandro Pecego Martins Romano, Pedro Paulo de Abreu Manso, Marcelo Pelajo-Machado, Patrícia Brasil, Cláudio Tadeu Daniel-Ribeiro, Cristiana Ferreira Alves de Britto, Maria de Fátima Ferreira-da-Cruz, Ricardo Lourenço-de-Oliveira

**Author notes:** Corresponding authors: MFFC; RLdO.

## Abstract

**Background:** Although malaria transmission was eradicated from southeast Brazil, a significant increase in the number of *Plasmodium vivax-like* autochthonous human cases has been reported in remote areas of the Atlantic Forest in the last decades in Rio de Janeiro (RJ) state, including an outbreak in 2015-2016. The singular clinical and epidemiological aspects of several human cases combined with molecular and genetic data revealed that they were due to the non-human primate (NHP) parasite *P. simium.* The full understanding of the epidemiology of the autochthonous malaria in southeastern Brazil depends, however, upon the knowledge on the circulation of NHP *Plasmodium* in the foci and the determination of its reservoirs.

**Methodology:** A large sampling effort was carried out in the Atlantic forest of RJ and its bordering states (Minas Gerais, São Paulo, Espírito Santo) for capture and examination of free-living NHPs. Blood and/or viscera were analyzed for Plasmodia infections through molecular and microscopic techniques.

**Principal findings:** In total, 146 NHPs of six species, from 30 counties in four states were tested. Howler monkeys (*A. guariba clamitans)* were the only NHP species found infected. In RJ, 26% of howlers were positive, among them 17% were found to be infected with *P. simium*. Importantly, specific single nucleotide polymorphisms were detected in all *P. simium* infected howlers regardless geographical origin of malaria foci. Interestingly, 71% of *P. simium* infected NHP were from the coastal slope of a mountain chain (Serra do Mar), where most human cases have been occurring. *P. brasilianum/malariae* was detected for the first time in 14% free-living howlers in RJ as well as in 25% of those from the Espírito Santo state. Moreover, malarial pigment was detected in spleen fragments of 50% of a subsample composed of howler monkeys found dead in both RJ and ES. All NHPs were negative for *P. falciparum*.

**Conclusions/Significance:** Our data indicate the howler monkeys as the main reservoir of the Atlantic forest human malaria in RJ and other sites in Southeast Brazil and reinforce its zoonotic nature.

**Author summary:** The present work consists of an unprecedented capture effort and large-scale field survey of plasmodial species in Non-human primates (NHPs) in RJ, a state recording a three-decade history of autochthonous human cases of benign tertian malaria pending epidemiological clarification of their origin. For the first time, we describe infection rates by *Plasmodium sp.*in free-living NHP, match the spatial distribution of *P. simium* in NHP with that of local human cases of benign tertian malaria due to this parasite, disclose howler monkeys as the only confirmed reservoir of this zoonotic malaria in the state and showed that specific single nucleotide polymorphisms were present in all *P. simium* infected howlers, regardless of the geographical origin of malaria foci. It is also the first time that *P. brasilianum/malariae* is recorded in free-living NHPs from Rio de Janeiro and the widespread distribution of this quartan-malaria parasite of zoonotic potential in the state is illustrated. Together, these findings increase the understanding about the simian malaria parasites in Atlantic Forests, as well as on the zoonotic character of autochthonous human malaria in Rio de Janeiro, providing subsidies for shaping surveillance and control.

## 1 Introduction

In Brazil, more than 99% of malaria infections are acquired in the Amazon, and few isolated imported cases or outbreaks of introduced cases from the Amazon or foreigner countries are occasionally recorded in the Extra-Amazonian regions [1]. Malaria transmission was considered eradicated from South and Southeast regions of Brazil more than 40 years ago [1]. However, in the last three decades, a significant increase in autochthonous malaria cases by *Plasmodium vivax-like* parasites in the Atlantic Forest areas in southeastern Brazil, where no index case that could have introduced the parasite from a malaria endemic region, have been reported [1,2]. These cases present similar parasitological, clinical and epidemiological characteristics, such as: low parasitemia, absence of the expected *P. vivax* relapses, and recent visits to areas covered by dense rain forest where the bromeliad-inhabiting *Anopheles* mosquitoes belonging to the subgenus *Kertezsia*, specially *An. cruzii,* are almost the exclusive human-biting anopheline species [2–4]. *An. cruzii* is the main vector of the so-called bromeliad malaria, endemic in Southern and Southeastern Brazil, and coincidently it is the only proved natural vector of simian malarias in the country [5]. This particular epidemiological context revived the hypothesis raised by Deane et al in the 1960’s about the existence of human malaria cases of simian origin in Brazil. In fact, these authors reported a human natural infection by the neotropical primate parasite *P. simium* Fonseca [6] in São Paulo (SP), southeast region [7]. The patient presented a benign tertian malaria after being exposed to mosquito bites at the tree-canopy during an entomological survey in a forest densely infested by *An. cruzii*. The description of vertical movement of *An. cruzii* between the canopy and ground level in the Atlantics rain forest of South Brazil reinforced Deane’s hypothesis that part of the transmission of bromeliad malaria in Southern and Southeastern Brazil would be of zoonotic character, being monkeys the parasite reservoir [5–8].

Two species of *Plasmodium* have been described in neotropical non-human primates (NHP): *P. brasilianum* Gonder e Berenberg-Gossler (1908) and *P. simium*, almost indistinguishable from the human malaria parasites *P. malariae* and *P. vivax*, respectively [5,6,9]. Besides subtle morphological variations [2,5], molecular markers such as microsatellites and single nucleotide polymorphisms (SNPs) were the only differences so far described between *P. malariae* and *P. brasilianum* and *P. vivax* and *P. simium* [2,10,11]. *P. brasilianum* has a broader distribution as has been found from México to Southern Brazil, infecting at least 11 genera of the five families of neotropical primates (Aotidae, Atelidae, Callitrichidae, Cebidae and Phiteciidae) [5,12–15]. In contrast, *P. simium* has been found essentially in species belonging to two genera (*Alouatta* and *Brachyteles*, family Atelidae) [5], from the Atlantic forest of Southern and Southeastern Brazilian. Importantly, both *P. brasilianum* and *P. simium* are experimentally infective to humans [9].

The full understanding of epidemiology of recent autochthonous malaria in southeastern Brazil depends on the confirmation of circulation of NHP Plasmodia in the transmission foci as well as the determination of parasite reservoirs. Studies on the prevalence of *P. simium* and/or *P. brasilianum* infections in NHPs and determine the potential reservoirs in southeastern Brazilian states were conducted during 1960-1990’s. Almost 800 NHPs were examined, recording a variation of *Plasmodium* infection from 10.9% in the states of Espírito Santo to 56.5% in SP [5]. At the time of these survey, only free-living lion-tamarins (Callitrichidae) could be examined from RJ and all were negative to malaria parasites [16]. However, RJ recorded 110 autochthonous human cases of benign tertian malaria between 2005 and 2018, with an outbreak in 2015-2016 of 49 cases [1,2]. Curiously, all these human infections were acquired in sites of RJ located along Serra do Mar, an extensive mountain chain covered by the best-preserved rain forest mosaic in the southeast. This biome harbor a rich NHP fauna, composed of species of six genera (*Alouatta*, *Brachyteles*, *Callicebus*, *Callithrix*, *Leontophitecus* and *Sapajus*) [17], and have *An. cruzi* as the most common anopheline [18]. Consequently, the hypothesis of simian origin in these RJ malaria cases has been raised [5,7]. In response, multidisciplinary studies including clinical, epidemiological, parasitological, and molecular approaches have been conducted in RJ [2,4,19]. More recently, molecular studies of parasites infecting humans and four howlers clearly demonstrated that they shared the same *P. simium* parasite. [2,7,20,21]. However, so far scarce number of wild NHPs of few species from only three out of numerous autochthonous malaria foci in RJ and surroundings could be examined [2]. This work presents the largest sampling effort ever carried out in the Atlantic forest of RJ and its borders for the capture and examination of free-living NHPs to describe the geographical distribution and frequency of simian malaria as well as to determine the local animal reservoirs and the identity of the parasite infecting humans and NHPs in the autochthonous malaria foci.

## 2 Material and Methods

### 2.1 Study area

The work was carried out between May 2015 and January 2019, totalizing approximately 120 days of fieldwork in 44 sites of 30 counties in the Atlantic Forest biome, in RJ as well as bordering sites in states of Minas Gerais (MG), ES and SP. The survey focused forest fragments from lowlands areas to mountain valleys and escarpments of Serra do Mar and other mountain chains [22]. The choice of capture areas comprised: local existence of NHPs and/or recent human malaria cases as well as alerts of an information network built with key institutions, inhabitants, health agents and environmental guards to continuously monitoring the presence of howler monkeys, as previously described [23].

### 2.2 Capture and sample collection

The expeditions consisted of ten-days surveys in the forests conducted by 2 - 6 trained people in target areas for the search of NHPs. The capture method was chosen according to NHP species, behavior and size [24]. Briefly, smallest and frugivorous species such as marmosets (genus *Callithrix*), lion tamarin (*Leontopithecus*), and capuchins (*Sapajus*) were trapped using banana baited automatic tomahawk traps [25]. Largest species presenting folivores and acrodendrophilic habits such as howlers (*Alouatta*) and woolly spider monkey (*Brachyteles*), or then which usually don’t enter in traps (e.g. one titi – *Callicebus*, captured in Minas Gerais) were captured with anesthetic darts, as previously described [23]. Sick animals reported by the information network during the 2017-2018 yellow fever epizooties [26] were captured with nets [23]. A sample of 3-6 mL of blood was collected from anesthetized or dying or recently dead animals. Thick and thin blood films were immediately prepared, and the remaining blood was let for coagulating. After sample collections and the complete recovery of anesthetic effects, animals were released always in the capture point during the daytime. Liver samples were obtained from recently dead animals, due to yellow fever or other disease. Liver and blood samples were stored at − 80°C until DNA extraction.

### 2.3 Malaria diagnosis

Giemsa-stained thick and thin blood films were examined in the microscopy under a x100 oil-immersion objective by two trained and independent microscopists. DNA was extracted from blood clots as previously described [27]; extractions from liver samples [28] were done using the QIAGEN DNeasy mini kit according to manufacturer’s instructions. Molecular diagnosis was made through conventional PCR. All DNA samples were tested in triplicate for 18s rRNA *Plasmodium* genus-specifc gene [29,30], and then for cysteine proteinase *P. vivax* and ssrRNA *P. malariae* and *P. falciparum* genes, as previously described [30,31]. *P. vivax-*positive samples were submitted to *P. simium* differential diagnosis based on a mitochondrial SNP [2,10]. The molecular diagnosis was performed by *Nested*-PCR of *coxI* gene fragment and subsequent enzymatic digestion, using primers and protocol previously described [10]. All the PCR products were visualized under UV light after electrophoresis on 2% agarose gels.

### 2.4 Histopathological analysis

Spleen fragments of a subsample consisting of 16 howlers (12 from RJ and 4 from ES) found dead were fixed in Carson’s formalin-Millonig [32] and processed according to standard histological techniques for paraffin embedding. Sections (5 μm thick) from each block were stained with hematoxylin-eosin [33] or Lennert Giemsa [34], and analyzed looking for malarial pigments under an AxioHome microscope equipped with an HRc Axiocam digital camera (Carl Zeiss, Germany).

### 2.5 Ethic issues

The collection methods, biosafety and anesthesia protocols adhered to the Brazilian law (11.794 of July 8, 2008) about the use of animals in scientific research, and complied with the rules and regulations of Brazilian Ministry of Health [24] having been previously approved by the institutional Ethics Committee for Animal Experimentation of Instituto Oswaldo Cruz (protocol CEUA/IOC-004/2015, license L-037/2016) and by Brazilian Ministry of the Environment (SISBIO 41837-3 and 54707-4) and Rio de Janeiro’s Environment agency (INEA 012/2016 and 019/2018). The research also adhered to the American Society of Primatologists Principles for the Ethical Treatment of Nonhuman Primates.

## 3 Results

In total, 146 animals belonging to six species from 30 counties in four Brazilian states were examined (S1 Table, Fig.1), being 130 by microscopy and PCR in blood samples, seven by microscopy and PCR in blood samples and histopathology of spleen fragments, and nine by PCR of viscera and histopathology of spleen fragments.

**Figure 1:**
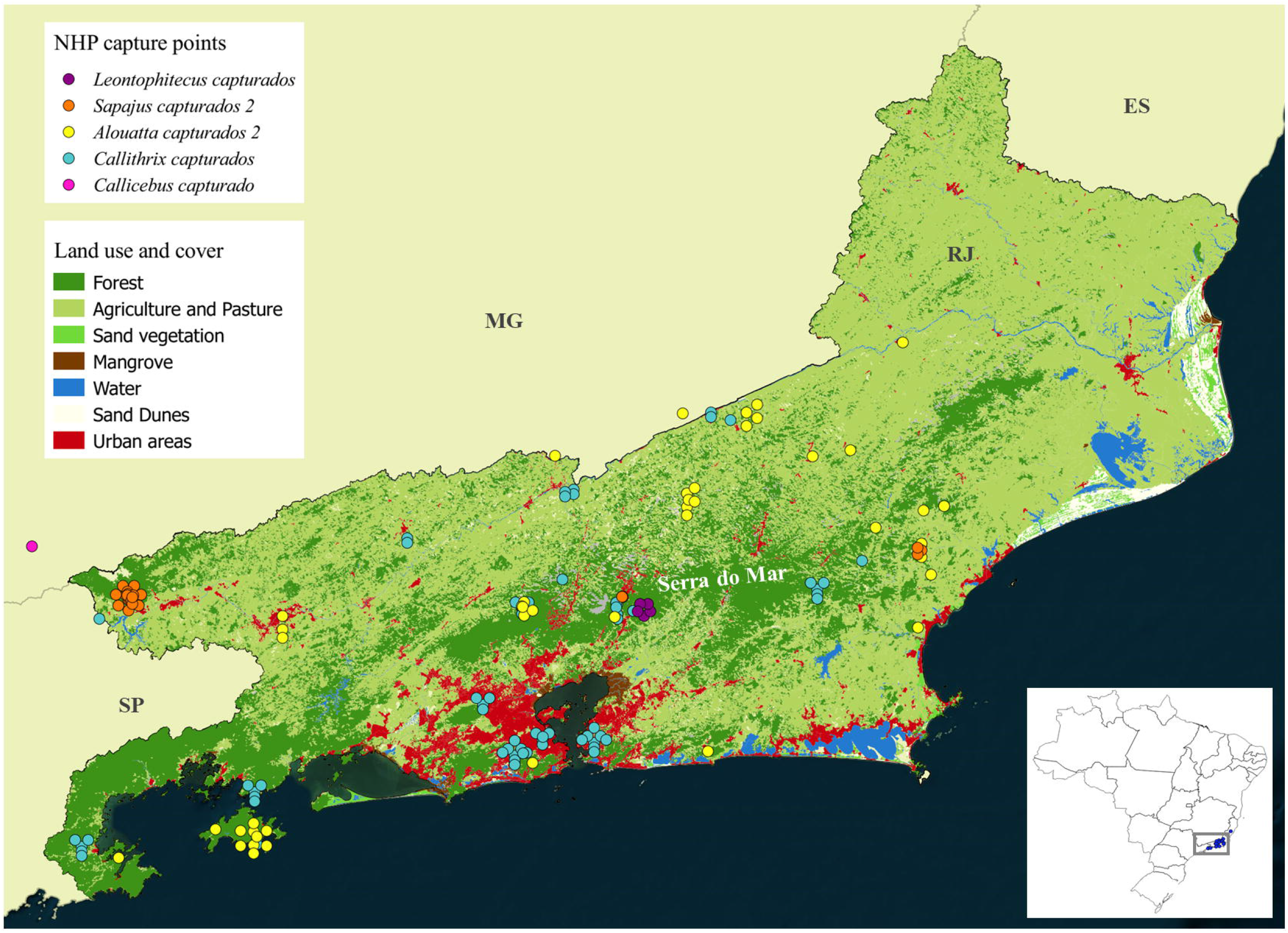
Map showing collection points of non-human primate in Rio de Janeiro. Each circle means one NHP examined. The map was prepared using free software QGIS 2.18.

Regardless the geographical origin, the only NHP species found infected with *Plasmodium* was the howler monkey *Alouatta g. clamitans* (Table 1). As expected, PCR was more sensitive than microscopic examination of blood films, that failed to detect *Plasmodium* in two infected howlers, one harboring *P. simium* and another *P. brasilianum/malariae* (Table 1). These results suggest that infected howler usually display detectable parasitemia despite the infecting plasmodial species. Trophozoites were the most common visualized blood form, but schizonts and gametocytes were also detected (Fig. 2). In addition, using PCR we were able to detect both *Plasmodium* species in four animals from which only liver and spleen samples were available.

**Table 1:** *Alouatta g. clamitans* captured and examined per state, infection rate per *Plasmodium* species and method of detection of current and past infection. Number (%).

| State | N | Total with *Plasmodium* | *P. simium* | *P. brasilianum* / *malariae* | *P. simium* and *P. brasilianum* / *malariae* | *P. falciparum* | Diagnosis |  | Malarial Pigment<sup>§</sup> |  |
| --- | --- | --- | --- | --- | --- | --- | --- | --- | --- | --- |
|  |  |  |  |  |  |  | Blood slides + PCR | Only PCR | N | Positive |
| **RJ** | 42 | 11 (26.1) | 5 (45.4) | 4 (36.3) | 2 (18.1) | - | 7<sup>#</sup> | 4<sup>*</sup> | 12 | 6 (50)<sup>1</sup> |
| **ES** | 4 | 1 (25.0) | - | 1 (100.0) | - | - | NR | 1 | 4 | 2 (50)<sup>2</sup> |
| **MG** | 2 | - | - | - | - | - | - | - | - | - |
| **TOT.** | **48** | **12 (25.0)** | **5 (41.6)** | **5 (41.6)** | **2 (16.6)** | **0** | **7** | **5** | **16** | **8 (50)<sup>3</sup>** |

<sup>#</sup>Parasitemia ranging from 15 to 300 parasites/μL (median = 40 p/μL); <sup>\*</sup>One harboring *P. simium*, one with *P. malariae/brasilianum* and the remaining two found dead with no blood slides available. NR: not realized, they were found dead during a yellow fever outbreak. <sup>§</sup>Search in spleen tissues in a subsample consisting of dead animals. <sup>1</sup>Two out of these six animals had positive PCR; <sup>2</sup>One of them had positive PCR. <sup>3</sup>Three of them had positive PCR; the other five had negative PCR.

**Figure 2:**
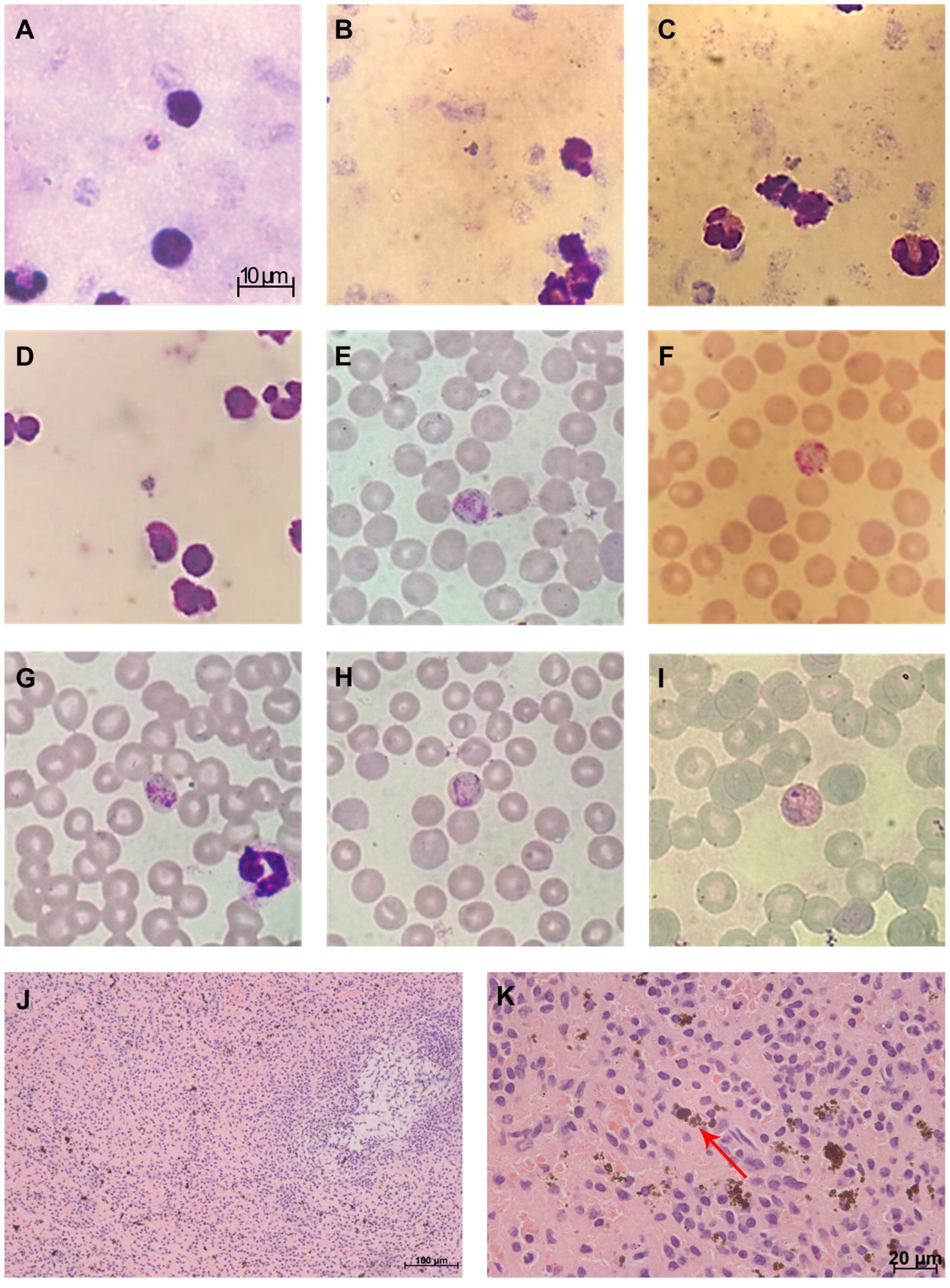
Giemsa’s solution-stained thick (A-D) and thin (E-I) blood samples, and histopathological analysis hematoxylin-eosin-stained spleen fragments of howlers naturally infected with *Plasmodium* in Rio de Janeiro state, Brazil, showing (J) hypertrophy of red pulp with malarial pigments and white pulp atrophy and (K) detail of malarial pigments in the red pulp.

Only 12 NHPs could be examined from the bordering states of RJ: six of them were howler monkeys from MG and ES and none was infected by *P. simium.* One of the four examined howlers from ES was PCR positive to *P. brasilianum/malariae* (25%) (Table 1).

Regarding RJ, 11 (26.1%) howler monkeys were infected with malaria parasites at the time of sampling and, among them, 16.7% were infected by *P. simium,* the causative agent of the autochthonous human malaria in this state (Tables 1 and 2). Importantly, specific *P. simium* SNP were detected in 100% of these tertian malaria parasite infected howlers. Coincidently, most of these animals were originated from the coastal slope of Serra do Mar where most human cases were recorded in the last decade (Table 2, Fig.3). Six howlers from RJ were infected by the quartan-malaria parasite *P. brasilianum/malariae* (14.3%), two of which displaying mixed *P. simium* infection (Tables 1 and 2). All samples were negative for *P. falciparum* parasites.

**Figure 3:**
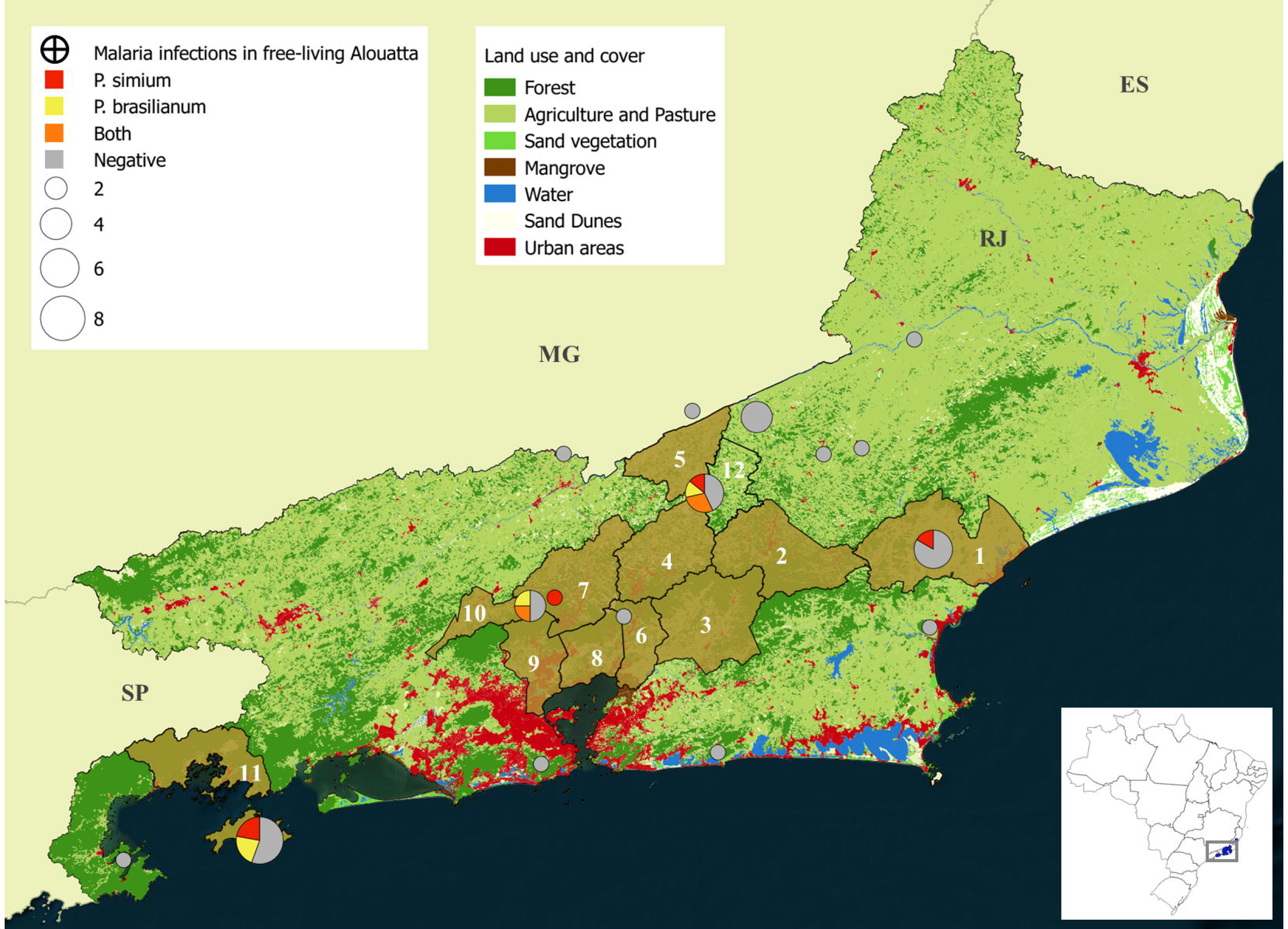
Map showing the distribution, number and *Plasmodium infections* of the examined *Alouatta g. clamitans* in Rio de Janeiro. Green shaded areas represent the counties with register of autochthonous malaria in human: 1. Macaé, 2. Nova Friburgo, 3. Cachoeira de Macacu, 4. Teresópolis, 5. Sapucaia, 6. Guapimirim, 7. Petrópolis, 8. Magé, 9. Duque de Caxias 10. Miguel Pereira, 11. Angra dos Reis. Number 12 represents Sumidouro, where human case has been never detected. The map was prepared using free software QGIS 2.18.

**Table 2:**
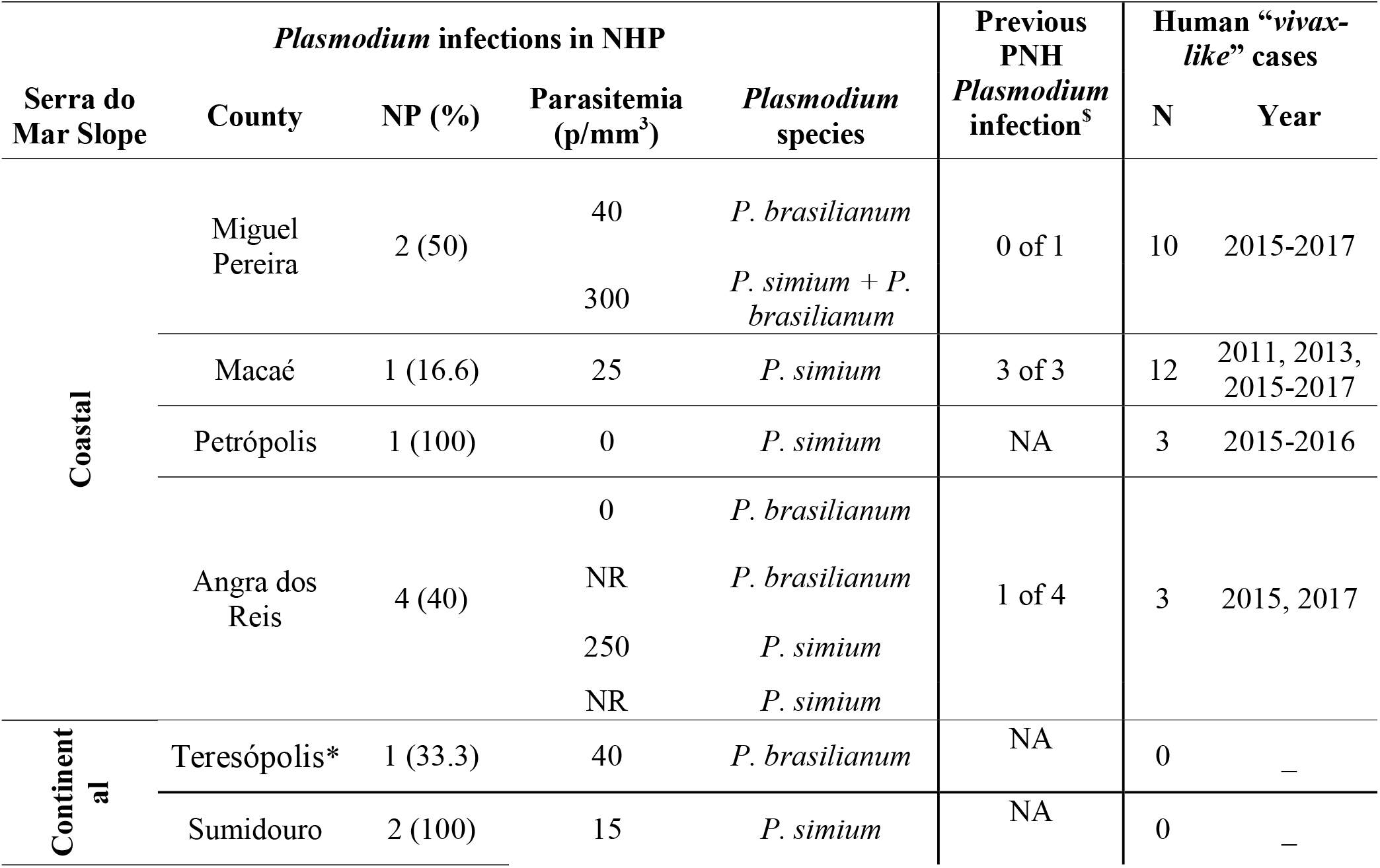
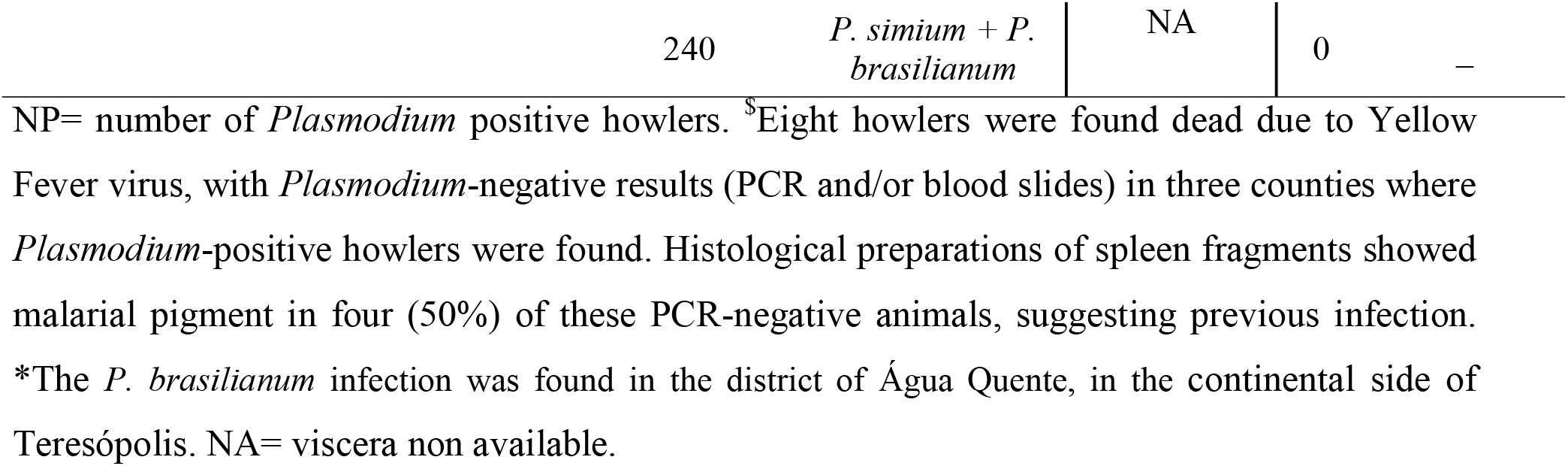
RJ *Plasmodium*-positive howler monkeys, according to plasmodial species, county, year, slope of capture, and occurrence of autochthonous human cases of benign tertian malaria recorded in the respective county and the year of detection.

Previous malaria infections could be investigated by the search of malarial pigment in a subsample of 16 howlers found dead. Accordingly, malarial pigment (Fig. 2) was detected in spleen fragments of five out of 13 animals with negative PCR at the time of death and, as expected, in three with positive PCR (Table 1). Interestingly, malarial pigment was found in spleen samples of 50% of howlers found dead in both RJ (6/12) and ES (2/4) displayed malarial pigment indicating that malaria is frequent in these monkeys from both states.

Howlers could be examined from 15 counties in RJ, six of which with records of autochthonous human malaria in the past decade. Current infections by *P. simium* were diagnosed in howlers from four (66.6%) of the surveyed counties reporting human cases of benign tertian malaria in the state, and in a neighboring county (Sumidouro) where human case has been never detected (Fig. 3). Coincidently, in Macaé where the highest number of human cases was recorded, all howlers found dead had malarial pigments in the spleen, suggesting that simian malaria is highly enzootic in the county (Table 2).

## 4 Discussion

The present work consists of an unprecedented capture effort and large-scale field survey of plasmodial species in NHPs in RJ, a state recording a three-decade history of autochthonous human cases of benign tertian malaria. For the first time, infection rates by *Plasmodium* were described, spatial distribution of *P. simium* in NHP was matched with local human cases of *P. simiun* malaria previously recorded, howler monkeys were disclosed as the only confirmed reservoir of this zoonotic malaria in the state and the presence of the so-called specific SNP was demonstrated in all *P. simium* infected howlers, regardless geographical origin of malaria foci. Although *P. brasilianum/malariae* has already been found in captive NHPs from RJ [13], it is also the first time that this parasite species was recorded in free-living NHPs from Rio de Janeiro and the widespread distribution of this quartan-malaria parasite and its zoonotic potential in the state were illustrated.

*P. simium* was detected in howlers captured in five out of 11 counties recently reporting autochthonous human malaria cases in RJ (Fig.2). Despite efforts, we failed in capturing howlers in four malaria foci due to some local hindrances such as the steep terrain, hunt pressure and low *A. g. clamitans* population densities [23]. Nevertheless, the strong geographical overlap of howler monkey and human infections by parasites displaying specific *P. simium* SNPs in 83.3% (five out of six) of malaria foci, strengthens the importance of howlers as main reservoir of benign tertian human malaria over the zoonotic transmission areas in Southeast Brazil [4,5,20,21]. Howlers have also been by far the NHP most commonly found parasitized with both *P. simium* and *P. brasilianum* in Southern and Southeastern Brazil [5]. Beside their susceptibility to *Plasmodium* infection [5], the acrodendrophilic behavior, the larger body surface, and the slower displacement (compared to the smaller monkeys) [35] may make them more exposed to the bites of the mosquito vectors. Importantly, the finding of malarial pigment in spleen fragments of 50% of a sub-sample consisting of howlers from RJ and ES, some of which *Plasmodium*-negative by PCR at the time of death, suggest that simian malaria is very frequent in this species. Indeed, in some areas with the highest numbers of human cases (e.g. Macaé), the percentage of howlers exposed to *Plasmodium sp* reached 66.6% when considering those with current and past malaria infections. This finding may also suggests that spontaneous healing from malaria infections may occur in howlers in nature, as described in *P. simium* experimental infections in some neotropical NHPs [36] and in a human natural infection [7].

The frequency of *P. simium* infection in free-living howler monkeys in RJ (16.6%) was higher than that found in the bordering state of SP (5.8%) but lower than that found in the entire southern and southeastern Brazilian regions (26.3-35%) [5,20,37]. *A. g. clamitans* was also the only free-living NHP from RJ in which blood forms by microscopy and plasmodial DNA were detected. Recently, DNA but not blood forms of *P. simium* was detected in captive *Cebus* and *Sapajus* from the Southeast, whose role as reservoir for the zoonotic malaria in the region is still unclear [28]. *P. brasilianum* DNA was detected in captive capuchin, titi, howler and owl monkeys, besides tamarins and marmosets [13,28], most of which was exotic species to the Brazilian Southeast. All these NHPs were confined in a breeding institution (Center of Primatology of Rio de Janeiro - CPRJ) located in a cleared area of the enzootic simian malaria forest in RJ. Therefore, it was suspected that the local ecological conditions favored the accidental infection of these captive NHPs by parasites carried from infected free-living howlers. One free living specimen was also found infected by *P. simium* near CPRJ [28]. There still no evidence if the parasite DNA found in the blood of these captive animals implies that they really undergo *Plasmodium* infections or only bear a transient parasitemia. However, it is important to continuously monitor the potential role as zoonotic plasmodia reservoir of other local or introduced NHPs, besides howlers.

Although *P. brasilianum* has been found in several NHP genera [5,38,39] around other Brazilian regions, previous studies conducted in Brazilian Atlantic forest and Cerrado biomes did not find any capuchin (56 examined) nor marmosets (out of 44) infected with *Plasmodium* [20]. In the same way, more than 270 marmosets and lion-tamarins from the Southeast were *Plasmodium* negative [5]. Splenectomized capuchins did not become infected when injected with *P. simium* infected blood, while splenectomized marmoset endured low parasitemia [36]. Thus, the epidemiological role of other NHPs besides howlers in the zoonotic transmission of malaria in Southeast Brazil, including RJ state, is probably despicable.

*P. brasilianum / malariae* was found in similar frequency to *P. simium* (14.3 versus 16.7%, respectively) in howlers from RJ, and mixed infections were recorded in 18% of *Plasmodium* infected ones. *P. brasilianum / malariae* was the only malaria parasite detected in howlers from ES (Table 1). Besides, geographical co-occurrence of these parasites seems to be frequent in RJ, as it was disclosed in three out of five counties wherein howlers were detected with malaria parasites (Fig. 2). Curiously, despite this coincident distribution and similar frequency of *P. brasilianum / malariae* and *P. simium* in RJ, autochthonous human cases in this and in other Atlantic forest states of Southeast Brazil (SP and ES) have been diagnosed microscopically and/or molecularly as benign tertian malaria due to *P. simium* for decades [1,4,5,7,21,40,41]. In reality, *P. simium* was only identified by molecular tests and DNAmt sequencing as the causative agent in the 2015-2016 malaria outbreak in RJ, whose patients were essentially non-residents of foci [2]. Nevertheless, six human asymptomatic infection by *P. malariae* were detected by PCR in residents of Guapimirim in RJ, in 2011 [19], and a subsample of reactive local individuals for any plasmodial species revealed antibodies against erythrocytic antigens of *P. malariae* in 30.9%. The hypothesis of infection of NHP origin due to *P. brasilianum* was raised because there was no index case of introduced or imported human case of *P. malariae* in Guapimirim, and because the cases had close contact with the Atlantic forest [19]. It is well known that *P. brasilianum* is a widespread and common simian malaria parasite in the Amazon [5,14,42] and that it is experimentally infective to humans, either by inoculation of parasitized monkey blood or by the bite of infected mosquitoes [9]. High prevalence of antibodies against sporozoites antigens and erythrocytic forms of *P. brasilianum/malariae* in people living or frequently working in Amazon forests (e.g. indians, miners, settlers) of Brazil, French Guiana and Venezuela suggested infection of this simian quartan malaria parasite to humans [42–44]. Infections by *P. brasilianum/malariae* in humans would be, therefore, expected to occur also in RJ and in other southeastern states where *P. simium* has been described. In effect, the natural vector of both parasites is the same (*An. cruzii*) [5]. Notwithstanding, why malaria cases due to *P. brasilianum/malariae* have not been reported yet in this region is a question that remains to clarify and it is recommended to strengthen malaria surveillance either in residents or visitors of the Atlantic forest to evaluate the zoonotic potential of *P. brasilianum/malariae* in southern and southeast Brazil [1].

Noteworthy, most of the *P. simium* - and *P. brasilianum/malariae* - infected howlers (73%) were from forest coastal slope of Serra do Mar, where all autochthonous human malaria cases have been acquired [2,19]. At least two main premises may explain this apparently geographical coincidence and distinct distribution of simian and human malarias. From the entomological and climatic points of view, the higher relative humidity in the costal slope may increase *An. cruzii* survival rates, supporting the sporogonic cycle of the *Plasmodia.* Sea moisture also favor density of epiphyte shade bromeliads, the larval habitat of *An. cruzii,* and generates higher rainfall indexes [45], which in turn increases the amount of water accumulated in the vector larval habitats, positively influencing mosquito density. Greater longevity and density directly influence the vector capacity of the mosquito to transmit *Plasmodia* [46–48]. Vector competence is governed by genetics of vector population and therefore influence *Plasmodium* transmission dynamics [48,49]. Indeed, Deane (1992) has emphasized that environmental conditions influence to a high degree the presence and densities of neotropical NHP hosts, bromeliads and *An. cruzii,* and, consequently, defining the occurrence or not of simian malaria in nearby sites. Moreover, two genetic lineages of *An. cruzii* with partial reproductive isolation have been recently described in Serra do Mar, one curiously occurring in the coastal and another in the continental slopes [50]. Costal slope of Serra do Mar has much more sites coveted by urban people to settle hidden weekend home in the forest and visited sights by ecotourists and hikers than the continental one. As explained, the autochthonous human cases in the Atlantic forest in RJ has been reported mainly in non-residents [1,2,19]. Together, the environmental, entomological, ecological and epidemiological characteristics seem to indicate costal slope of Serra do Mar as the place to acquire malaria of simian origin. Protective measures such as the use of repellents and long clothes should be encouraged especially for those who live or practice ecotourism in this slope.

During the present study, an yellow fever virus (YFV) outbreak erupted in the southeast Brazil, a region without records of this virus circulation for almost 80 years [26,51]. Hundreds of epizootics of NHPs were reported, causing a significant impact on the population size of howler monkeys, an extremely susceptible host to YFV [24,52–57]. Considering the role of the howlers as reservoir of *Plasmodium* infective to humans, it is plausible to suppose that dynamics of zoonotic transmission of malaria will undergo changes in short or medium term in RJ and bordering states affected by the YFV epizooties. In this context, we postulate that the rapid decrease of *Alouatta* populations would also decrease the source of plasmodial infection to *An. cruzii*, which would reduce the circulation of *Plasmodia* in the Atlantic forest. Despite the short time since the 2016-2018 YFV epizootics, records from the malaria surveillance seem to confirm this scenario. In fact, there was an abrupt drop in human malaria case records between 2018-2019 [58], contrasting with the numbers reported in 2015-2016 [2]. If the reduction of autochthonous malaria cases in Atlantic Forest is depending of plasmodial sources of NHP origin the role of NHP for the occurrence of malaria in Extra-Amazonia would be reinforced as never before.

Previous sampling efforts of examining free-living NHPs in the Southeastern Atlantic Forest over the last 30 years showed limited geographical coverage, with samplings essentially limited to wildlife rescues or carried out in areas close to cities, or were based on few individuals [4,20,21]. As a result, our data contribute to understanding the simian malaria parasite distribution and frequency as well as the zoonotic character of autochthonous human malaria in Rio de Janeiro, which in turn provides subsidies for shaping surveillance and control. The evidence of the simian origin of parasites infecting humans and the widespread occurrence of anophelines vectors in the southeast increased the concern of the reemergence of endemic or epidemic autochthonous transmission in the region independent of the enzootic cycle [2]. However, it is not clear if parasitaemic humans infected by the bite of *An. cruzii* carrying esporozoites of *P. simium* acquired from howlers could be source of infection to *An. cruzii* or any other malaria vector occurring in the region. It is knowing that *P. simium* infected humans usually display scanty to null parasitemia, can cure spontaneous in few days without treatment and any relapse or molecular detection of parasites during follow-ups of treatment have been described [2,5]. Besides, all autochthonous human case of benign tertian malaria detected for decades in southeast have reported recent contact with the *P. simium* enzootic forest, and no secondary transmission directly derived from a human infected in the zoonotic cycle has never been detected outside the sylvatic foci. These epidemiological and parasitological profiles appear to indicate that humans are not a source of *P. simium* infection to mosquitoes. In this light, determining vector competence of *An. cruzii* and other traditional human malaria vector occurring in the southeast (*e.g*. *An. darlingi, An. aquasalis* and *An. albitarsis*) for transmiting *P. simium* and *P. brasilianum* between humans and from NHPs and humans and vice-versa is imperative.

## Authors’ contributions

RLO Conceptualization; CFAB, CTDR, MFFC, RLO Funding acquisition APMR, APC, CBJ, DAMA, DST, ES, FVSA, LRG, MPM, MQG, PPAM, RLO, WPV Investigation; APC, CBJ, CFAB, DAMA, FVSA, MQG, MFFC, PB, RLO Data curation; APC, FVSA, PB, RLO Formal analyses; APMR, ES, CFAB, DAMA, DST, FVSA, MFFC, RLO Methodology; FVSA, RLO Project administration; APMR, CTDR, CFAB,MPM, MFFC, RLO Resources FVSA, PPAM, MPM, RLO Visualization; FVSA, MFFC, RLO Writing original draft; APC, CFAB, CTDR, DAMA, DST - Review and Editing the manuscript; All authors read and approved the manuscript.

## Acknowledgments

To Carlos Alberto C. da Silva, Alexandre B. de Souza, Vicente Klonowski, Romenique L. Araújo, Luiz R. Nogueira, Fernando Barreto, Ana L. Quijada, Luiz P. P. Silva, Gelson Medeiros, Adilson B. Ramos, Marcilene B. Ramos, Carlos A. A. Júnior, Paulo G. Barbosa, Sérgio F. Fragoso, Adilson R. Silva, Cecília Cronemberger, Marcelo Rheingantz, Leonardo Nascimento, João Marins for the support in the field. To Orzinete Rodrigues Soares for non◻human primates’ blood slides’ review. To Grupo Técnico de Vigilância de Arboviroses (GT◻Arbo—Secretaria de Vigilância em Saúde—Brazilian Ministry of Health) for field and material supports.

## Supplementary table 1

Number of examined NHP, by species, habitat and capture method.

## References

1. Pina-Costa A de, Brasil P, Santi SM Di, Araujo MP de, Suárez-Mutis MC, Santelli ACF e S, et al. Malaria in Brazil: what happens outside the Amazonian endemic region. Mem Inst Oswaldo Cruz. 2014;109: 618–633. doi:10.1590/0074-0276140228

2. Brasil P, Zalis MG, de Pina-Costa A, Siqueira AM, Júnior CB, Silva S, et al. Outbreak of human malaria caused by *Plasmodium simium* in the Atlantic Forest in Rio de Janeiro: a molecular epidemiological investigation. Lancet Glob Heal. 2017;5. doi:10.1016/S2214-109X(17)30333-9

3. Curado I, dos Santos Malafronte R, de Castro Duarte AMR, Kirchgatter K, Branquinho MS, Bianchi Galati EA. Malaria epidemiology in low-endemicity areas of the Atlantic Forest in the Vale do Ribeira, São Paulo, Brazil. Acta Trop. 2006;100: 54–62. doi:10.1016/j.actatropica.2006.09.010

4. Cerutti C, Boulos M, Coutinho AF, Hatab M do CL, Falqueto A, Rezende HR, et al. Epidemiologic aspects of the malaria transmission cycle in an area of very low incidence in Brazil. Malar J. 2007;6: 33. doi:10.1186/1475-2875-6-33

5. Deane LM. Simian malaria in Brazil. Mem Inst Oswaldo Cruz. 1992;87: 1–20. doi:10.1590/S0074-02761992000700001

6. Fonseca F. Plasmódio de primata do Brasil. Mem Inst Oswaldo Cruz. 1951. pp. 543–553. doi:10.1590/S0074-02761951000100008

7. Deane LM, Deane MP, Ferreira Neto J. Studies on transmission of simian malaria and on a natural infection of man with *Plasmodium simium* in Brazil. Bull World Health Organ. 1966;35: 805–8.

8. Deane LM, Ferreira Neto JA, Lima MM, Deane LM, Ferreira Neto JA, Lima MM. The vertical dispersión of *Anopheles* (*Kerteszia*) *cruzi* in a forest in southern Brazil suggests that human cases of malaria of simian origin might be expected. Mem Inst Oswaldo Cruz. 1984;79: 461–463. doi:10.1590/S0074-02761984000400011

9. Coatney GR. The Simian Malaria: Zoonoses, Anthroponoses, or Both? Am J Trop Med Hyg. 1971;20: 795–803.

10. de Alvarenga DAM, Culleton R, de Pina-Costa A, Rodrigues DF, Bianco C, Silva S, et al. An assay for the identification of *Plasmodium simium* infection for diagnosis of zoonotic malaria in the Brazilian Atlantic Forest. Sci Rep. 2018;8: 86. doi:10.1038/s41598-017-18216-x

11. Guimarães LO, Bajay MM, Wunderlich G, Bueno MG, Röhe F, Catão-Dias JL, et al. The genetic diversity of *Plasmodium malariae* and *Plasmodium brasilianum* from human, simian and mosquito hosts in Brazil. Acta Trop. 2012;124: 27–32. doi:10.1016/j.actatropica.2012.05.016

12. Rylands AB, Mittermeier RA, Silva JS. Neotropical primates: taxonomy and recently described species and subspecies. Int Zoo Yearb. 2012;46: 11–24. doi:10.1111/j.1748-1090.2011.00152.x

13. Alvarenga DAM, Pina-Costa A, Bianco C, Moreira SB, Brasil P, Pissinatti A, et al. New potential *Plasmodium brasilianum* hosts: tamarin and marmoset monkeys (family Callitrichidae). Malar J. 2017;16: 71. doi:10.1186/s12936-017-1724-0

14. Lourenço-de-Oliveira R, Deane LM. Simian malaria at two sites in the Brazilian Amazon. I--The infection rates of *Plasmodium brasilianum* in non-human primates. Mem Inst Oswaldo Cruz. 1995. pp. 331–339. doi:10.1590/S0074-02761995000300004

15. Eyles DE. The species of simian malaria: Taxonomy, morphology, life cycle, and geographical distribution of the monkey species. J Parasitol. 1963;49: 866–87. doi:10.2307/3275712

16. Lourenço-de-Oliveira R, Ziccardi M. Natural infection of Golden lion tamarin, *Leontopithecus rosalia*, with *Trypanosoma cruzi*, in the state of Rio de Janeiro, Brazil. Mem Inst Oswaldo Cruz. 1990;85: 15.

17. Paglia AP, Rylands AB, Herrmann G, Aguiar LMS, Chiarello AG, Leite YLR, et al. Lista anotada dos mamiferos do Brasil, segunda edicao. 2nd ed. Arlington: Conservation International; 2012.

18. Guimarães AE, Arlé M, Machado RNM. Mosquitos no Parque Nacional da Serra dos Órgãos, Estado do Rio de Janeiro, Brasil. II. Distribuição vertical. Mem Inst Oswaldo Cruz. 1985;80: 171–85.

19. Miguel RB, Albuquerque HG, Sanchez MCA, Coura JR, Santos S da S, Silva S da, et al. Asymptomatic *Plasmodium* infection in a residual malaria transmission area in the Atlantic Forest region: Implications for elimination. Rev Soc Bras Med Trop. 2019;52: 1–9. doi:10.1590/0037-8682-0537-2018

20. de Castro Duarte AMR, Malafronte R dos S, Cerutti C, Curado I, de Paiva BR, Maeda AY, et al. Natural *Plasmodium* infections in Brazilian wild monkeys: Reservoirs for human infections? Acta Trop. 2008;107: 179–185. doi:10.1016/j.actatropica.2008.05.020

21. Buery JC, Rodrigues PT, Natal L, Salla LC, Loss AC, Vicente CR, et al. Mitochondrial genome of *Plasmodium vivax/simium* detected in an endemic region for malaria in the Atlantic Forest of Espírito Santo state, Brazil: do mosquitoes, simians and humans harbour the same parasite? Malar J. 2017;16: 437. doi:10.1186/s12936-017-2080-9

22. Ribeiro MC, Metzger JP, Martensen AC, Ponzoni FJ, Hirota MM. The Brazilian Atlantic Forest: How much is left, and how is the remaining forest distributed? Implications for conservation. Biol Conserv. 2009;142: 1141–1153. doi:10.1016/J.BIOCON.2009.02.021

23. Abreu FVS, dos Santos E, Gomes MQ, Vargas WP, Oliveira Passos PH, Nunes e Silva C, et al. Capture of *Alouatta guariba clamitans* for the surveillance of sylvatic yellow fever and zoonotic malaria: Which is the best strategy in the tropical Atlantic Forest? Am J Primatol. 2019;81: e23000. doi:10.1002/ajp.23000

24. Brasil. Guia de vigilância de epizootias em primatas não humanos e entomologia aplicada à vigilância da febre amarela. 2nd ed. Brasília: Ministério da Saúde; 2017.

25. Watsa M, Erkenswick G, Halloran D, Kane EK, Poirier A, Klonoski K, et al. A Field Protocol for the Capture and Release of Callitrichids. Neotrop Primates. 2015;22: 59–68.

26. Possas C, Lourenço-de-oliveira R, Tauil PL, Pinheiro FDP, Pissinatti A, Venâncio R, et al. Yellow fever outbreak in Brazil◻: the puzzle of rapid viral spread and challenges for immunisation. 2018;113: 1–12. doi:10.1590/0074-02760180278

27. de Abreu FVS, Gomes LR, Mello ARL, Bianco-Júnior C, de Pina-Costa A, dos Santos E, et al. Frozen blood clots can be used for the diagnosis of distinct *Plasmodium* species in man and non-human primates from the Brazilian Atlantic Forest. Malar J. 2018;17: 338. doi:10.1186/s12936-018-2485-0

28. de Alvarenga D, de Pina-Costa A, de Sousa T, Pissinatti A, Zalis MG, Suaréz-Mutis MC, et al. Simian malaria in the Brazilian Atlantic forest: first description of natural infection of capuchin monkeys (Cebinae subfamily) by *Plasmodium simium*. Malar J. 2015;14: 81. doi:10.1186/s12936-015-0606-6

29. Gama BE, Silva-Pires F do ES, Lopes MNR, Cardoso MAB, Britto C, Torres KL, et al. Real-time PCR versus conventional PCR for malaria parasite detection in low-grade parasitemia. Exp Parasitol. 2007;116: 427–432. doi:10.1016/j.exppara.2007.02.011

30. Torres KL, Figueiredo D V., Zalis MG, Daniel-Ribeiro CT, Alecrim W, Ferreira-da-Cruz M de F. Standardization of a very specific and sensitive single PCR for detection of *Plasmodium vivax* in low parasitized individuals and its usefulness for screening blood donors. Parasitol Res. 2006;98: 519–524. doi:10.1007/s00436-005-0085-8

31. Snounou G, Viriyakosol S, Xin Ping Zhu, Jarra W, Pinheiro L, do Rosario VE, et al. High sensitivity of detection of human malaria parasites by the use of nested polymerase chain reaction. Mol Biochem Parasitol. 1993;61: 315–320. doi:10.1016/0166-6851(93)90077-B

32. Carson FL, Martin JH, Lynn JA. Formalin Fixation for Electron Microscopy: A Re-evaluation. Am J Clin Pathol. 1973;59: 365–373. doi:10.1093/ajcp/59.3.365

33. Mayer P. Die Caprellidae der Siboga-Expedition. Leiden, Buchhandlung und druckerei vormals E.J. Brill; 1903.

34. Lennette DA. An improved mounting medium for immunofluorescence microscopy. Am J Clin Pathol. 1978;69: 647–648. doi:10.1093/ajcp/69.6.647

35. Kowalewski MM, Garber PA, Cortés-Ortiz L, Urbani B, Youlatos D. Howler monkeys◻: behavior, ecology, and conservation. New York, NY: Springer; 2014. doi:10.1007/978-1-4939-1960-4

36. Deane LM. Studies on simian malaria in Brazil. Bull World Health Organ. 1964;31: 752–3.

37. Costa DC, Cunha VP da, Assis GMP de, Souza Junior JC de, Hirano ZMB, Arruda ME de, et al. *Plasmodium simium/Plasmodium vivax* infections in southern brown howler monkeys from the Atlantic Forest. Mem Inst Oswaldo Cruz. 2014;109: 641–653. doi:10.1590/0074-0276130578

38. Lourenço-de-Oliveira R, Luz SLB. Simian malaria at two sites in the Brazilian Amazon - II. Vertical distribution and frequency of anopheline species inside and outside the forest. Mem Inst Oswaldo Cruz. 1996;91: 687–694. doi:10.1590/S0074-02761996000600005

39. Figueiredo MAP, Di Santi SM, Manrique WG, André MR, Machado RZ. Identification of *Plasmodium* spp. in Neotropical primates of Maranhense Amazon in Northeast Brazil. PLoS One. 2017;12: 1–14. doi:10.1371/journal.pone.0182905

40. Gomes ADC, De Paula MB, Duarte AMRDC, Lima MA, Malafronte RDS, Mucci LF, et al. Epidemiological and ecological aspects related to malaria in the area of influence of the lake at Porto Primavera dam, in western São Paulo State, Brazil. Rev Inst Med Trop Sao Paulo. 2008;50: 287–295. doi:10.1590/S0036-46652008000500008

41. Brasil P, Costa A, Longo C, Silva S, Ferreira-da-Cruz M, Daniel-Ribeiro CT. Malaria, a difficult diagnosis in a febrile patient with sub-microscopic parasitaemia and polyclonal lymphocyte activation outside the endemic region, in Brazil. Malar J. 2013;12: 402.

42. de Arruda M, Nardin EH, Nussenzweig RS, Cochrane AH. Sero-epidemiological studies of malaria in Indian tribes and monkeys of the Amazon Basin of Brazil. Am J Trop Med Hyg. 1989;41: 379–85.

43. Volney B, Pouliquen J-F, De Thoisy B, Fandeur T. A sero-epidemiological study of malaria in human and monkey populations in French Guiana. Acta Trop. 2002;82: 11–23. doi:10.1016/S0001-706X(02)00036-0

44. Lalremruata A, Magris M, Vivas-Martínez S, Koehler M, Esen M, Kempaiah P, et al. Natural infection of *Plasmodium brasilianum* in humans: Man and monkey share quartan malaria parasites in the Venezuelan Amazon. EBioMedicine. 2015;2: 1186–1192. doi:10.1016/j.ebiom.2015.07.033

45. Brito TT, Oliveira-Júnior JF, Lyra GB, Gois G, Zeri M. Multivariate analysis applied to monthly rainfall over Rio de Janeiro state, Brazil. Meteorol Atmos Phys. 2017;129: 469–478. doi:10.1007/s00703-016-0481-x

46. Machado RL, Ceddia MB, Carvalho DF de, Cruz ES da, Francelino MR. Spatial variability of maximum annual daily rain under different return periods at the Rio de Janeiro state, Brazil. Bragantia. 2010;69: 77–84. doi:10.1590/S0006-87052010000500009

47. Coutinho JO, Rachou R. Data on the biology and the malaria vector capacity of anophelines of sub-group *Kerteszia* under natural conditions. Rev Bras Malariol Doencas Trop. 1966;18: 557–80.

48. Cohuet A, Harris C, Robert V, Fontenille D. Evolutionary forces on *Anopheles*: what makes a malaria vector? Trends Parasitol. 2010;26: 130–136. doi:10.1016/J.PT.2009.12.001

49. Beerntsen BT, James AA, Christensen BM. Genetics of mosquito vector competence. Microbiol Mol Biol Rev. 2000;64: 115–37. doi:10.1128/mmbr.64.1.115-137.2000

50. de Rezende Dias G, Fujii TTS, Fogel BF, Lourenço-de-Oliveira R, Silva-do-Nascimento TF, Pitaluga AN, et al. Cryptic diversity in an Atlantic Forest malaria vector from the mountains of South-East Brazil. Parasit Vectors. 2018;11: 36. doi:10.1186/s13071-018-2615-0

51. Bonaldo MC, Gómez MM, dos Santos AA, Abreu FVS de, Ferreira-de-Brito A, Miranda RM de, et al. Genome analysis of yellow fever virus of the ongoing outbreak in Brazil reveals polymorphisms. Mem Inst Oswaldo Cruz. 2017;112: 447–451. doi:10.1590/0074-02760170134

52. Brasil. Monitoramento do Período Sazonal da Febre Amarela [Internet]. 2018 pp. 1–15. Available: http://portalarquivos2.saude.gov.br/images/pdf/2018/fevereiro/21/Informe-n14-FA-20fev18-c.pdf

53. Moreno ES, Agostini I, Holzmann I, Di Bitetti MS, Oklander LI, Kowalewski MM, et al. Yellow fever impact on brown howler monkeys (*Alouatta guariba clamitans*) in Argentina: A metamodelling approach based on population viability analysis and epidemiological dynamics. Mem Inst Oswaldo Cruz. 2015;110: 865–876. doi:10.1590/0074-02760150075

54. Almeida M., Dos Santos E, Da Cruz Cardoso J, Da Fonseca DF, Noll CA, Silveira VR, et al. Yellow fever outbreak affecting *Alouatta* populations in southern Brazil (Rio Grande do Sul State), 2008-2009. Am J Primatol. 2012;74: 68–76. doi:10.1002/ajp.21010

55. Araújo FAA, Ramos DG, Santos AL, Passos PH de O, Elkhoury ANSM, Costa ZGA, et al. Epizootias em primatas não humanos durante reemergência do vírus da febre amarela no Brasil, 2007 a 2009. Epidemiol e Serviços Saúde. 2011;20: 527–536. doi:10.5123/S1679-49742011000400012

56. Bicca-marques JC, Calegaro-marques C, Rylands AB, Karen B, Mittermeier RA, Almeida MAB De, et al. Yellow fever threatens Atlantic Forest primates Júlio. Sci Adv. 2017;1600946: 18–20.

57. Duchiade A. Brazilian forests fall silent as yellow fever decimates threatened monkeys. In: Scientific American [Internet]. 12 May 2018 [cited 12 May 2019]. Available: https://www.scientificamerican.com/article/brazilian-forests-fall-silent-as-yellow-fever-decimates-threatened-monkeys/

58. Datasus Ministério da Saúde. TabNet Win32 3.0: MALÁRIA - Casos confirmados Notificados no Sistema de Informação de Agravos de Notificação - Rio de Janeiro [Internet]. [cited 12 May 2019]. Available: http://tabnet.datasus.gov.br/cgi/tabcgi.exe?sinannet/cnv/malarj.def

