## Supplementary material for "Howler monkeys are the reservoir of malaria parasites causing zoonotic infections in the Atlantic forest of Rio de Janeiro": S1 Table

Supplementary table 1: Number of examined NHP, by species, habitat and capture method.

| **Species** | **N** | **Brazilian states** | | | | **Capture method** | | | |
| --- | --- | --- | --- | --- | --- | --- | --- | --- | --- |
|  |  |  |  |  |  | N **(%)** | | | |
|  |  | **RJ** | **MG** | **SP** | **ES** | | **Darted Trapped Other^§^** | | |
| *Alouatta g. clamitans* | **48** | 42 | 2 | - | 4 | | 12 (25) | 0 (0) | 36 (75) |
| *Brachyteles arachnoides* | **1** | - | - | - | 1 | | 1 (100) | 0 (0) | 0 (0) |
| *Callicebus nigrifrons* | **1** | - | 1 | - | - | | 1 (100) | 0 (0) | 0 (0) |
| *Callithrix jacchus* and hybrids*** | **66** | 62 | 3 | 1 | - | | 0 (0) | 61 (93) | 5 (7) |
| *Leontopithecus rosalia* | **5** | 5 | - | - | - | | 0 (0) | 0 (0) | 5 (100) |
| *Sapajus nigritus* | **25** | 25 | - | - | - | | 0 (0) | 24 (96) | 1 (4) |
| **Total** | **146** | 134 | 6 | 1 | 5 | | 14 (9.6) | 85 (58.2) | 47 (32.2) |

*Most individuals presented phenotype of *Callithrix jacchus* x *Callithrix penicillate* and one individual was a *C. jacchus* x *Callithrix aurita* hybrid. ^§^Animals dying or recently dead, or captured with nets following alert from our information network.
